# MEDiCINe: Motion Correction for Neural Electrophysiology Recordings

**DOI:** 10.1101/2024.11.06.622160

**Authors:** Nicholas Watters, Alessio Buccino, Mehrdad Jazayeri

## Abstract

Electrophysiology recordings from the brain using laminar multielectrode arrays allow researchers to measure the activity of many neurons simultaneously. However, laminar microelectrode arrays move relative to their surrounding neural tissue for a variety of reasons, such as pulsation, changes in intracranial pressure, and decompression of neural tissue after insertion. Inferring and correcting for this motion stabilizes the recording and is critical to identify and track single neurons across time. Such motion correction is a preprocessing step of standard spike sorting methods. However, estimating motion robustly and accurately in electrophysiology recordings is challenging due to the stochasticity of the neural data. To tackle this problem, we introduce MEDiCINe (Motion Estimation by Distributional Contrastive Inference for Neurophysiology), a novel motion estimation method. We show that MEDiCINe outperforms existing motion estimation methods on an extensive suite of simulated neurophysiology recordings and leads to more accurate spike sorting. We also show that MEDiCINe correctly estimates the motion in primate electrophysiology recordings with a variety of motion and stability statistics. We open-source MEDiCINe, usage instructions, examples integrating MEDiCINe with common tools for spike-sorting, and data and code for reproducing our results. This open software will enable other researchers to use MEDiCINe to improve spike sorting results and get the most out of their electrophysiology datasets.

## 1. Introduction

Electrophysiology studies often involve recording neural activity with laminar microelectrode arrays inserted in the brain. This data is processed to compute putative spike times of individual neurons throughout a recording session, a process termed “spike sorting”. Historically, spike sorting was a primarily manual process. However, recent advances in recording scale afforded by high-density laminar microelectrode arrays such as Neuropixels probes (Jun et al., 2017) have made manual spike sorting prohibitively time-consuming. This has necessitated the emergence of automated spike sorting algorithms (Buccino et al., 2022). Automating spike sorting is challenging for several reasons, one of which is that the laminar microelectrode array (hereafter referred to as “array”) may move relative to its surrounding neural tissue (Figure 1-A) (Garcia et al., 2024; Steinmetz et al., 2018). This motion can be caused by a variety of factors, such as pulsation, changes in intracranial pressure, decompression of neural tissue after inserting an array, and instability of the mechanical apparatus holding the array. Motion is typically more extreme in non-human primate (NHP) and human recordings than recordings in rodents and other small animals (Coughlin et al., 2023; Windolf et al., 2023). Estimating and correcting for motion is an important step in spike-sorting pipelines. Improvements in motion estimation yield better automatic spike sorting, both yielding more usable neurons for analysis and saving the researcher time manually curating spike sorting results (Garcia et al., 2024).

**Figure 1.**
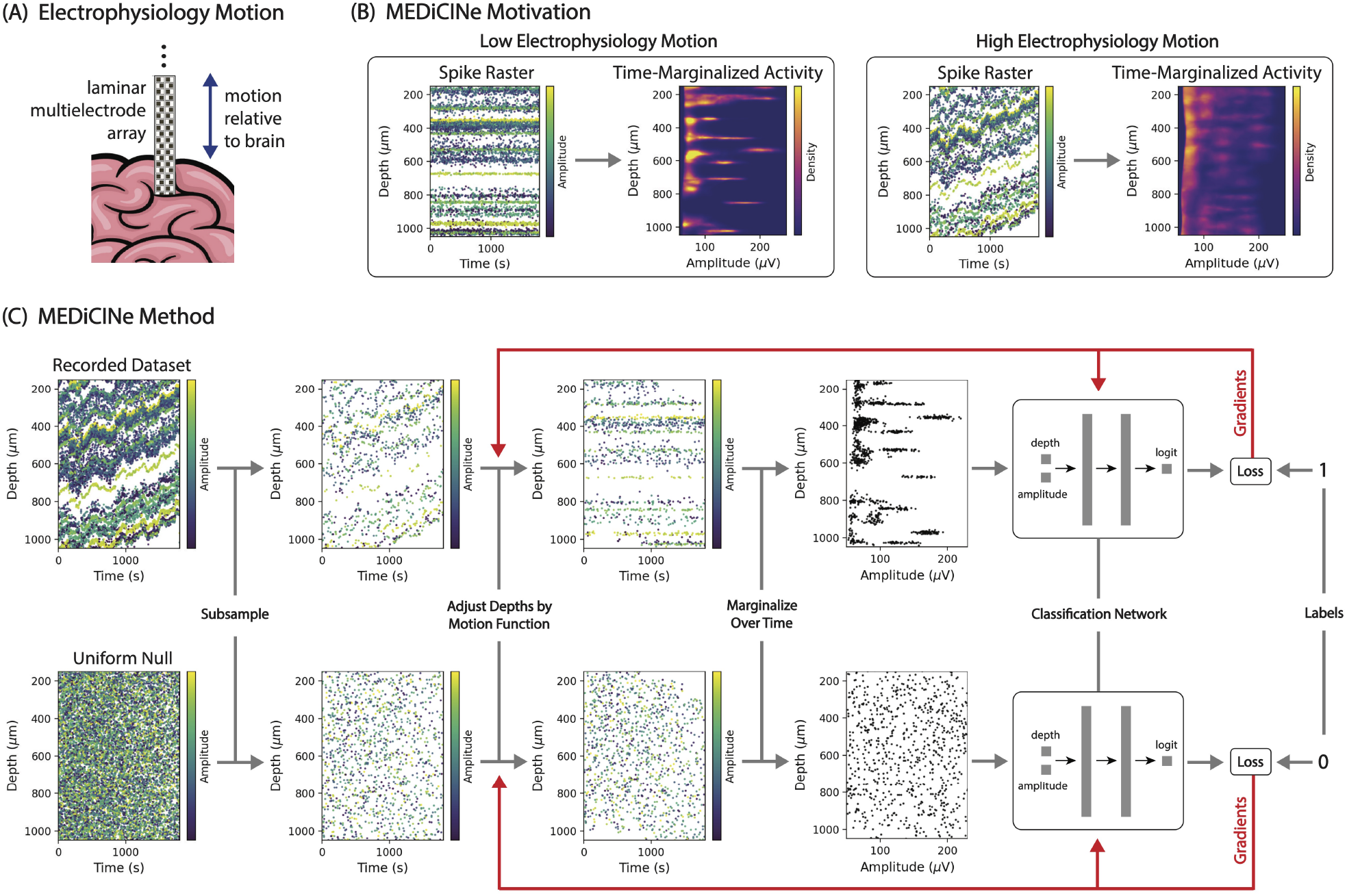
MEDiCINe is a method for estimating motion in electrophysiology data. **(A)** Laminar multielectrode array (checkerboard) inserted into brain (pink) moves relative to the brain over time in a recording session. **(B)** In recordings with low electrophysiology motion, neural activity does not vary in depth over time, causing timemarginalized activity distribution to be sparse. In recordings with high electrophysiology motion, neural activity varies in depth over time, causing time-marginalized activity distribution to not be sparse. **(C)** MEDiCINe motion-estimation procedure. For each training step, spikes are sampled from the recorded dataset and from a uniform null distribution. Each spike has a time, depth, and amplitude feature. Spike depths are adjusted by a differentiable motion function, then marginalized over time. A classification network then discriminates between spikes that came from the dataset and spikes that came from the uniform distribution. Binary cross-entropy classification loss is backpropagated to update the parameters of both the classification network and the motion function.

Electrophysiology motion estimation is challenging for several reasons. First, the motion can exhibit a range of statistics, including slow drift, high-frequency noise, and discrete jumps. Second, the motion may depend on position along the array, varying as a function of depth in the brain. Third, the neural activity itself may be non-stationary: Neuron firing rates may fluctuate over time, neurons may be gained or lost throughout the session due to motion or cell death, and the relative motion between the array and the brain may cause the recorded waveform shape of single neurons to change.

Existing state-of-the-art approaches to motion estimation begin by discretizing the data into temporal bins (Pachitariu et al., 2024; Windolf et al., 2023). For each temporal bin, they compute a histogram of neural activity as a function of depth along the array and neural activity features such as waveform amplitude. An estimate of the motion is computed to maximize the correlations across pairs of these histograms. This computation may involve comparing each temporal bin to a particular reference bin (Pachitariu et al., 2024), or may involve comparing each temporal bin only to its nearest neighbors (Windolf et al., 2023). While these approaches work well for some recordings, there is broad agreement in the field that existing approaches struggle for some recordings and accurate motion estimation is a common difficulty for spike sorting.

We introduce **MEDiCINe** (**M**otion **E**stimation by **Di**stributional **C**ontrastive **I**nference for **Ne**urophysiology), an approach for motion estimation that infers motion by fitting a constrained model of the neural data. We first consider the generative process of the neural data, which has two components: (1) Neural activity consisting of local voltage modulations, such as spikes or LFP power, coming from neurons that are unmoving in the brain tissue, and (2) Motion of the array relative to the brain. We then formulate a non-parametric model of neural activity that captures this generative structure. We fit this model to neural data using gradient descent. We found that this approach works on a wide range of datasets without hyperparameter tuning. MEDiCINe out-performs existing methods on an extensive suite of simulated datasets with known ground-truth motion and a variety of motion and instability statistics. MEDiCINe also works well on all of our NHP Neuropixels recordings. Lastly, we open-source MEDiCINe, usage instructions, examples integrating MEDiCINe with SpikeInterface and Kilosort4 tools for spike sorting, and data and code for reproducing our results.

## 2. Methods

Consider a dataset of *N* spikes (putative action potentials) extracted from an electrophysiology recording session. Represent the dataset as a set of 3-vectors [(*t*_1_, *d*_1_, *a*_1_), (*t*_2_, *d*_2_, *a*_2_), …, (*t*_*N*_, *d*_*N*_, *a*_*N*_)] where *t*_*i*_ is the time at which spike *i* occurred, *d*_*i*_ is the estimated depth along the laminar array at which spike *i* was detected, and *a*_*i*_ is the amplitude of spike *i*. If the recording has low electrophysiology motion through time, then marginalizing this dataset over time would yield a sparse distribution in depth-amplitude space, assuming individual neurons have stable depth and spike amplitude in the brain. In contrast, if the recording has high motion, then marginalizing this dataset over time would not yield a sparse dataset, because spikes coming from a single neuron would be spread out over depth (Figure 1-B). Leveraging this observation, MEDiCINe learns a motion function to maximize sparsity of the time-marginalized dataset distribution (Figure 1-C).

To operationalize this, MEDiCINe learns two differentiable functions:

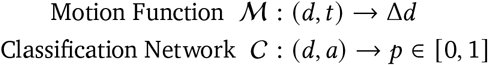

These functions compose to form a probability function 𝒫 over the joint space [time, depth, amplitude]:

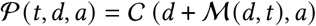

ℳ and 𝒞 are fit using gradient descent. For each step, we draw a batch 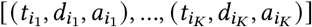 of *K* random samples from the spike dataset and a batch 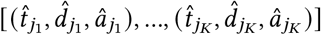 of *K* random samples from a uniform distribution with the same ranges at the spike dataset. We then apply 𝒫 to each of these batches to get 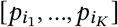 and 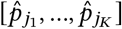, where 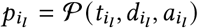 and 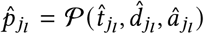. We then compute the loss function

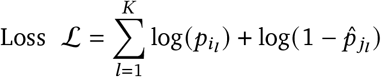

This is the binary cross entropy loss where the data samples are labeled 1 and the uniform samples are labeled 0. We backpropagate ℒ to update the parameters in ℳ and 𝒞. This loss function pressures 𝒫 to discriminate dataset samples from uniform samples, hence pressuring the motion function to cause the time-marginalized dataset distribution after motion adjustment to be sparse.

We parameterize the classification network 𝒞 by a multilayer perceptron with 2 hidden layers. We parameterize the motion function ℳ as the linear interpolation of a matrix of shape [depth bins, time bins] discretizing the space of depth and time. The entries of this matrix at Δ*d* estimates. Using multiple depth bins allows the motion function to model motion that varies across depth.

Note that this method is not specific to datasets with only [time, depth, amplitude] representations of spikes. It can apply more generally to any dataset of neural events that have a [time, depth, feature vector] representation. This includes spike data with spike shape features beyond amplitude and LFP data with power spectrum features. Note also that this method is not specific to motion only in depth. By letting the motion function return a 3-vector [Δ*x*, Δ*y*, Δ*z*], it could estimate motion in 3-dimensional space.

See Supplementary Section 5.2 for additional details and open-sourced code.

## 3. Results

To evaluate MEDiCINe and compare it to existing motion estimation methods, we quantitatively benchmarked it on a suite of simulated datasets with controlled ground-truth motion. We also qualitatively assessed its performance on NHP electrophysiology datasets without known ground-truth motion.

### 3.1. Simulated Datasets

To compare the performance of MEDiCINe with existing motion estimation methods, we generated a suite of 384 simulated neurophysiology recording datasets with controlled ground-truth motion and evaluated both MEDiCINe and existing motion estimation methods on these datasets. To generate the simulated datasets, we enlarged a preexisting suite of simulated datasets (Garcia et al., 2024) to include a wide variety of motion and neuron stability statistics that occur in neurophysiology data. We used the MEArec electrophysiology data simulator (Buccino and Einevoll, 2021) to generate one 30-minute Neuropixels electrophysiology session for each combination of the following dataset parameters:

- Linear drift of the relative depth between array and brain. Two options: (i) No linear drift or (ii) linear drift of 0.1*μm* / *s*.
- Random walk of the relative depth between array and brain with Gaussian steps and 1-second frequency. Three options: (i) No random walk, (ii) random walk with standard deviation 1 *μm* / *s*, or (iii) random walk with standard deviation 2 *μm* / *s*.
- Discrete random jumps of the relative depth between array and brain. Two options: (i) No jumps or (ii) jump times sampled from a Poisson process with rate 100*s* and jump displacements sampled from a uniform distribution over [−50*μm*, 50*μm*].
- Number of neurons. Two options: (i) 20 neurons, or (ii) 100 neurons.
- Distribution of neuron density over depth. Two options: (i) Neurons uniformly distributed over depth or (ii) neurons distributed bimodally over depth.
- Firing rate stability. Four options: (i) Constant firing rates, (ii) periodic firing rates that are synchronous over all neurons, (iii) periodic firing rates that are asynchronous over all neurons, or (iv) half of the neurons have constant firing rates while the other half appear or disappear at random times in the session with linearly ramping firing rate.
- Depth-dependency of motion. Two options: (i) depth-independent (rigid) motion or (ii) depth-dependent (non-rigid) motion that varies linearly over depth.

From these datasets we extracted spike times, estimated depths, and amplitudes using the monopolar triangulation method (Boussard et al., 2021; Buccino et al., 2018). See Figure 2-A for spike times from three example simulated datasets and Figure 2-B for results of MEDiCINe applied to these datasets.

**Figure 2.**
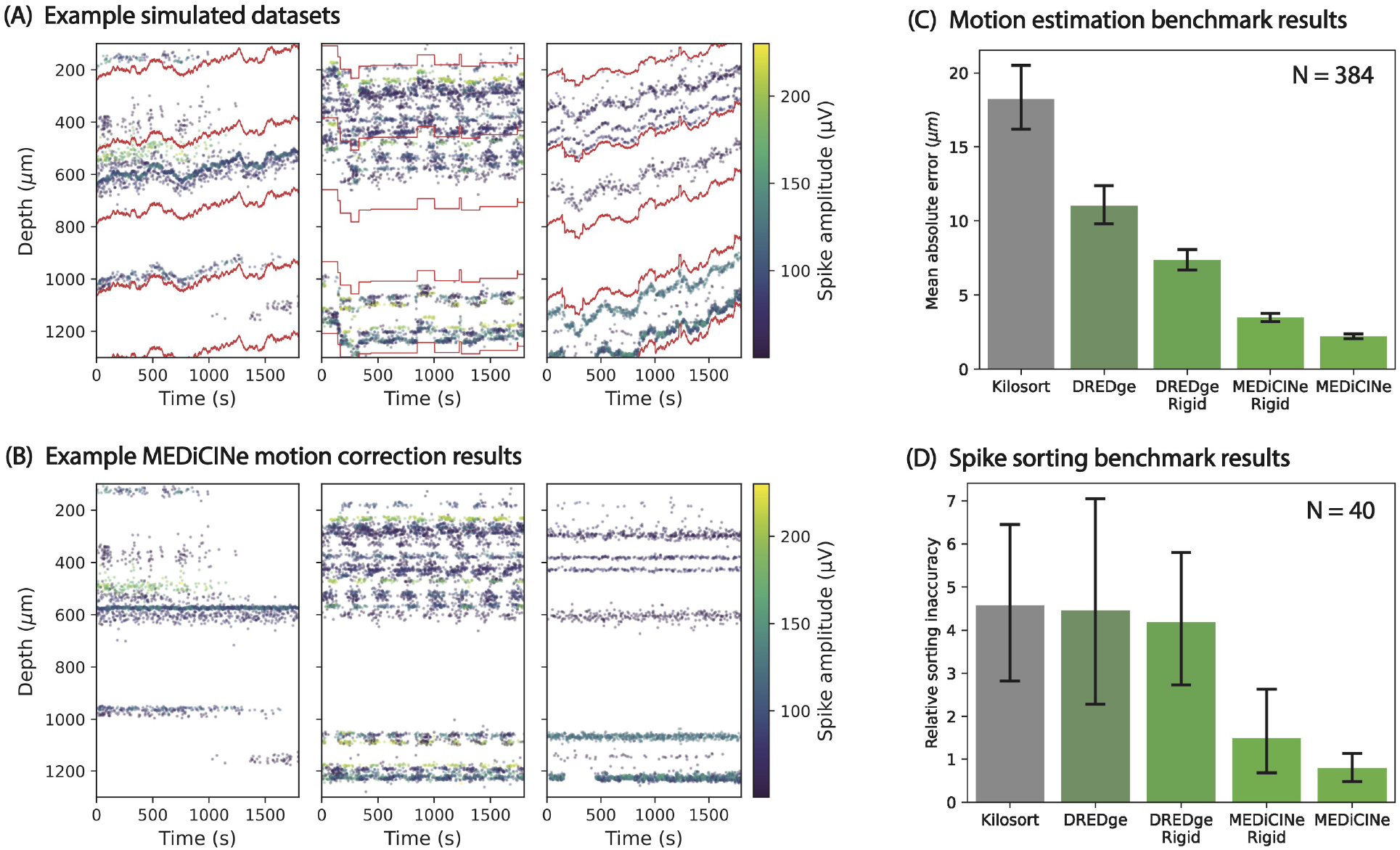
Simulated data results. **(A)** Examples for 3 of our 384 simulated datasets used for benchmarking. Each point represents a detected spike. Only 3% of spikes are shown. Red lines show the ground-truth motion, namely the depth on the array over time of stationary points in the brain. The left dataset has random walk motion with standard deviation 2 *μm* / *s*, 20 neurons, and firing rate instability consisting of appearing/disappearing neurons. The middle dataset has motion consisting of discrete random jumps, 100 neurons with bimodal density over depth, and asynchronous periodic firing rate instability. The right dataset has depthdependent motion consisting of both linear drift and random walk with standard deviation 2 *μm* / *s*, and 20 neurons with bimodal density over depth. **(B)** Spike rasters after correcting for motion estimated by MEDiCINe for the three example datasets in (A). **(C)** Mean absolute error (y-axis) between estimated motion and ground-truth motion over all 384 simulated datasets for each motion estimation method (x-axis). Errorbars are 95% confidence intervals of the mean. Kilosort = 18.23 ± 2.12, DREDge = 11.03 ± 1.32, DREDge Rigid = 7.36 ± 0.69, MEDiCINe Rigid = 3.48 ± 0.28, MEDiCINe = 2.21 ± 0.17. **(D)** Mean spike sorting inaccuracy (y-axis) over 40 simulated datasets between ground-truth spike times per unit and Kilosort4 estimated spike times per unit after motion estimation for each motion estimation method (x-axis). Errorbars are 95% confidence intervals of the mean. Kilosort = 4.57 ± 1.87, DREDge = 4.46 ± 2.33, DREDge Rigid = 4.19 ± 1.61, MEDiCINe Rigid = 1.50 ± 0.94, MEDiCINe = 0.79 ± 0.34.

We evaluated the following five motion estimation methods on the extracted spikes from each of our 384 simulated datasets:

- *Kilosort*. The “datashift” motion estimation function from Kilosort4 (Pachitariu et al., 2024), which is currently the most recent motion estimation in the Kilosort family.
- *DREDge*. The official DREDge implementation in the SpikeInterface library (Buccino et al., 2020), currently considered the state-of-the-art motion estimation method (Windolf et al., 2023).
- *DREDge Rigid*. A modification of DREDge that enforces rigid motion as a function of depth and uses center-of-mass depth estimation instead of monopolar triangulation (Garcia et al., 2024).
- *MEDiCINe Rigid*. Our MEDiCINe method with a single depth bin, enforcing rigid motion as a function of depth.
- *MEDiCINe*. Our MEDiCINe method with multiple depth bins. In practice we used 2 depth bins, which is the same number as DREDge uses on our simulated datasets with the default parameters.

We evaluated performance of these motion estimators by measuring the mean absolute error of estimated motion with respect to the ground-truth motion. MEDiCINe Rigid and MEDiCINe significantly outperformed all other methods on average (Figure 2-C). When conditioning these results on each factor of variation of the datasets, MEDiCINe always performed at least as well as all existing methods (Supplementary Figure 4). These results are not due to outlier effects (Supplementary Figure 5). On a per-dataset basis, MEDiCINe Rigid and MEDiCINe also ranked highest on average among all the methods (Supplementary Figure 6).

Prior work has shown that better motion estimation correlates with better spike sorting (Garcia et al., 2024). To verify this, we selected a random set of 40 of our simulated datasets to evaluate spike sorting. For each of these datasets and each motion estimation method, we corrected for the estimated motion in the neural data using Kriging interpolation (Pachitariu et al., 2023) and ran Kilosort4 spike sorting (disabling the built-in motion correction step) (Pachitariu et al., 2024). To evaluate sorting quality, we computed a standard metric of spike-sorting accuracy (Garcia et al., 2024; Pachitariu et al., 2024). We then define for any motion estimation method, the inaccuracy on a dataset to be the difference between the accuracy of that method and the accuracy of the most accurate method on the dataset. MEDiCINe Rigid and MEDiCINe had lower sorting inaccuracy than existing methods (Figure 2-D). See Supplementary Section 5.4.7 for methodological details.

### 3.2. Primate Datasets

To test MEDiCINe in practice, we used four of our primate Neuropixels sessions with motion artifacts that we found difficult to estimate and correct using existing methods. These recordings exhibited a range of real-world motion and instability conditions. We again used monopolar triangular spike localization and applied MEDiCINe. We found that MEDiCINe performed well under these conditions (Figure 3), qualitatively better than existing methods on these datasets (Supplementary Figure 9).

**Figure 3.**
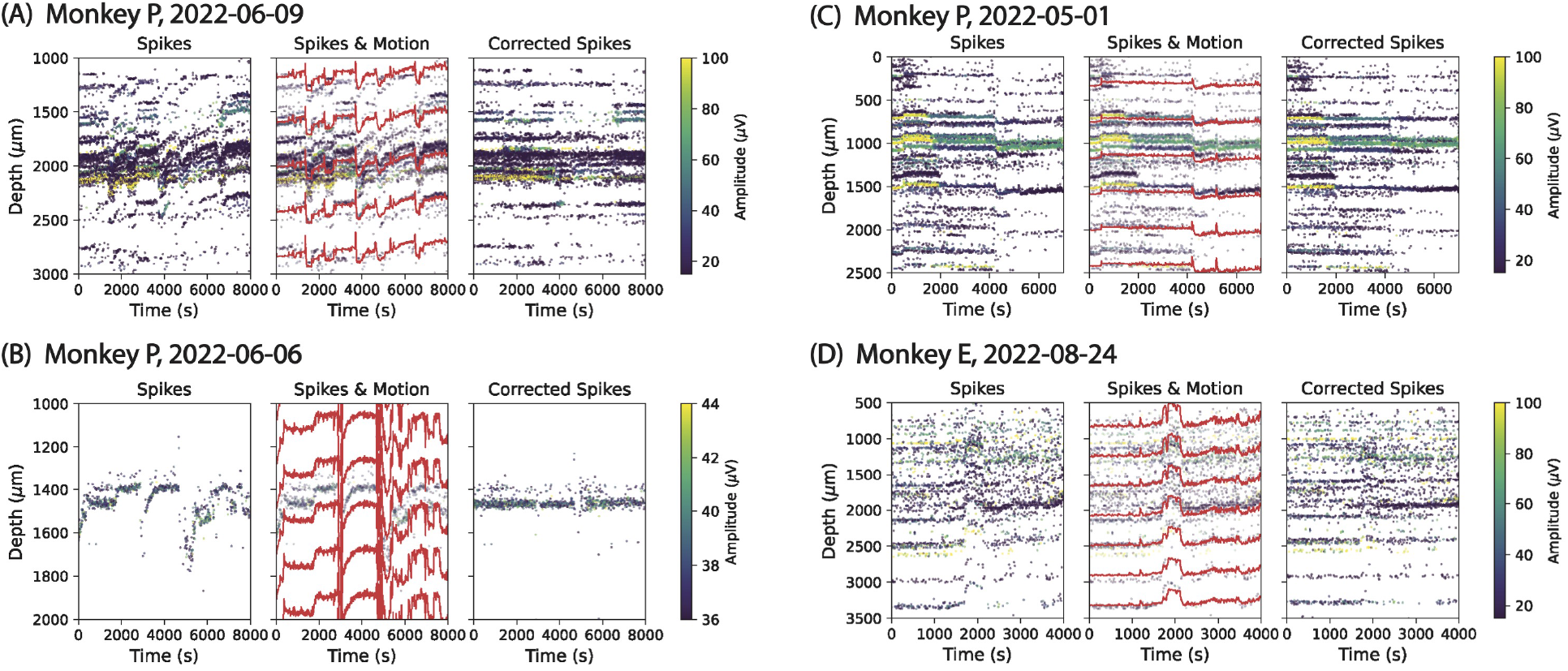
Primate data results. **(A) - (D)** Each panel represents a different NHP Neuropixels dataset that we recorded, showing a raster plot of spikes *(left)*, interpolated motion curves estimated by MEDiCINe (red lines) overlayed on the raster plot *(middle)*, and the raster plot after correcting by the motion estimated by MEDiCINe *(right)*.

**Figure 4.**
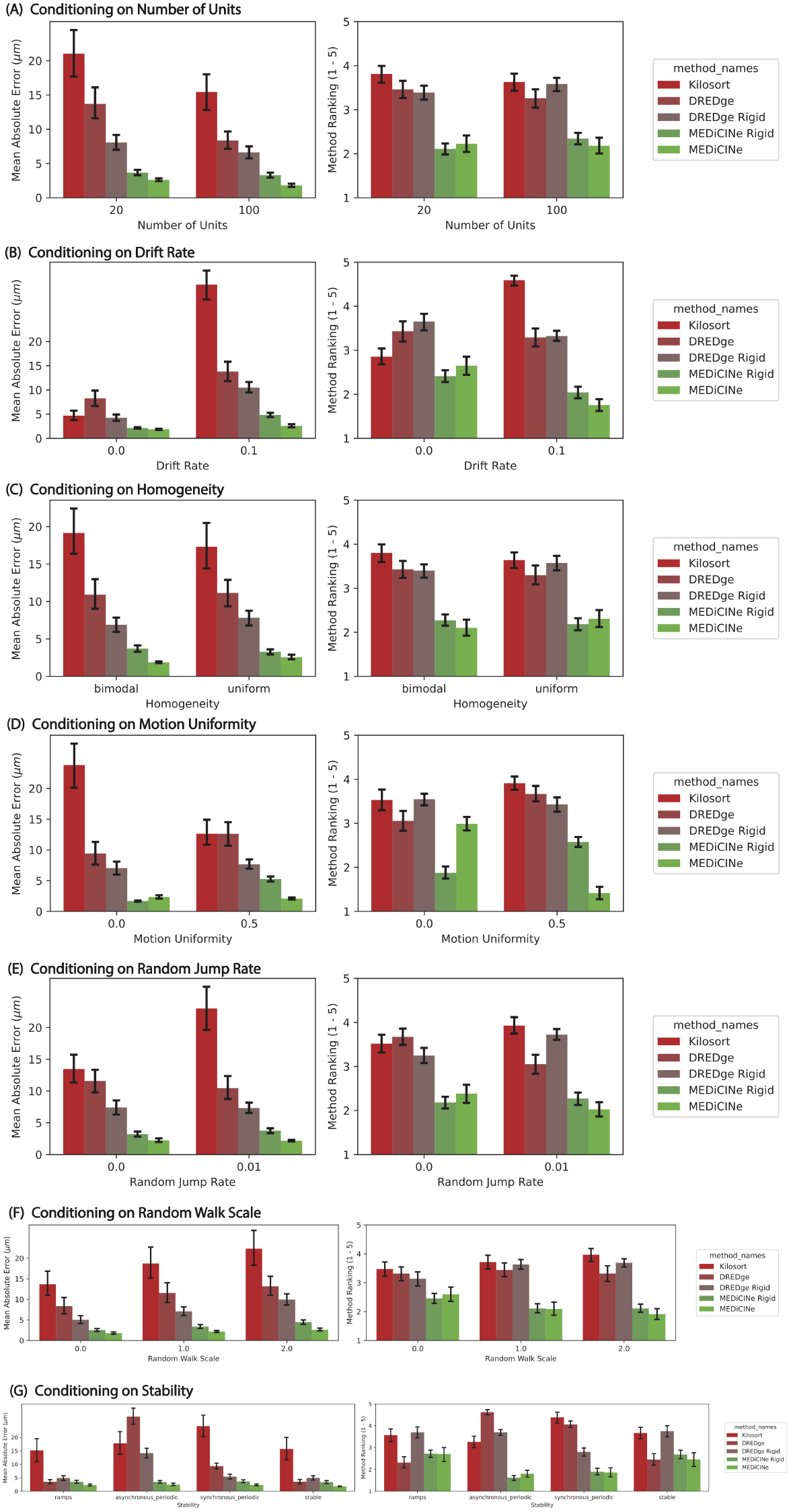
Results on Simulated Data Conditioned on Parameters. Motion estimation model results conditioned on each parameter of variation of simulated dataset suite. Errorbars show 95% confidence interval of the mean.

**Figure 5.**
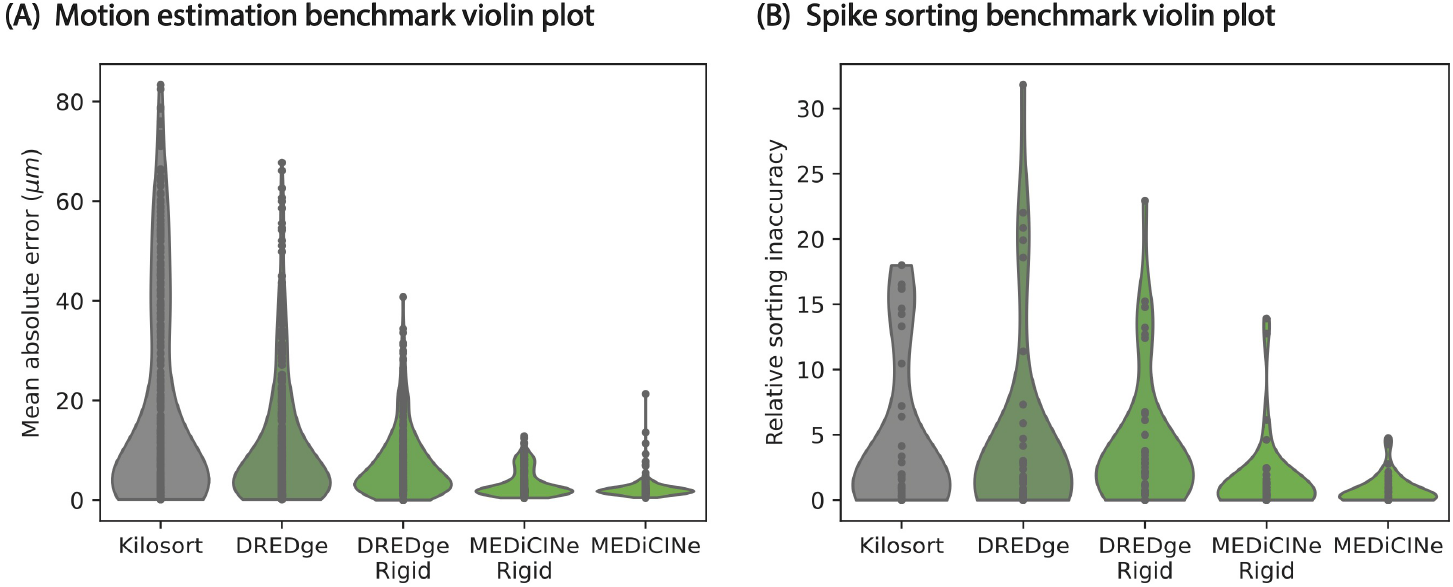
Benchmark Violin Plots. **(A)** A violin plot representation of the results in Figure 2-A. **(B)** A violin plot representation of the results in Figure 2-B.

**Figure 6.**
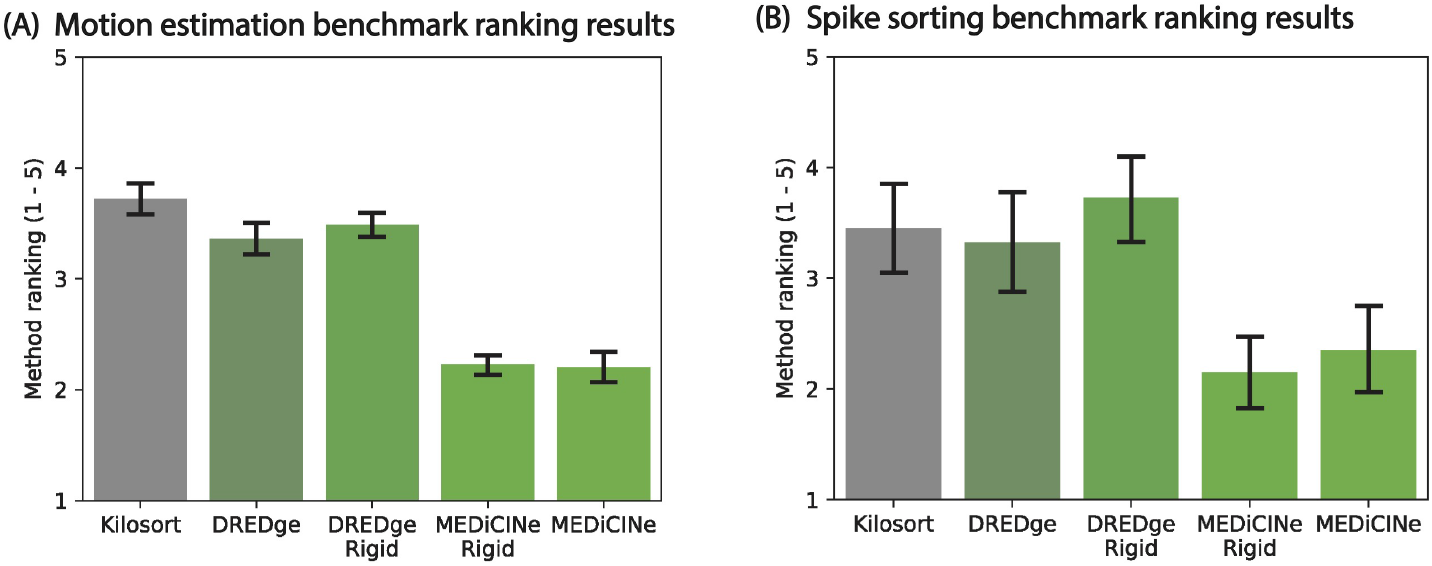
Benchmark method Rankings. **(A)** For each simulated dataset, we compute the ranking (1-5) of each of the 5 motion estimation methods on that dataset in terms of mean absolute motion estimation error. This ranking is shown on the y-axis. **(B)** For each simulated dataset for which we run spike sorting, we compute the ranking (1 - 5) of each of the 5 motion estimation method on that dataset in terms of relative sorting inaccuracy. This ranking is shown on the y-axis.

## 4. Discussion

In this work, we introduced a novel method for estimating motion in neurophysiology recordings, called MEDiCINe (Motion Estimation by Distributional Contrastive Inference for Neurophysiology). We found that MEDiCINe out-performed existing methods on an extensive benchmark of simulated datasets with known ground-truth motion. We also found that MEDiCINe performed well on real NHP neurophysiology datasets where existing methods struggle.

We envision several ways to extend and improve the MEDiCINe model:

- LFP features. In this work we have evaluated MEDiCINe on spike data, but using LFP data may allow it to estimate motion more accurately, particularly for datasets with few neurons. Other works have found LFP features useful for motion estimation (Windolf et al., 2023).
- More spike shape features. In this work the only spike shape feature we used for motion estimation was amplitude. However, MEDiCINe could readily use other features, such as spike width or waveform shape. In fact, because MEDiCINe uses a sparsity loss based on classification, we expect using more spike features would improve its performance by increasing the sparsity of the motion-corrected time-marginalized spike distribution.
- Fluctuations in firing rate over time. In this work, we used a time-invariant classification network 𝒞 to discriminate dataset spikes from uniformly sampled spikes. However, in practice neuron firing rates change over time (e.g. due to cell death). Modeling these changes in firing rate could improve the performance of MEDiCINe. One way to do this would be to allow the classification network to depend on time subject to reasonable priors, such as only allowing sparse or slow changes in firing rates.
- Inductive biases on motion. In this work, the motion function *M* was unconstrained aside from a temporal smoothing kernel. This affords the model flexibility, but causes it to sometimes find implausible solutions (Supplementary Figure 7). This could be addressed by incorporating more priors in the motion function, such as a Gaussian process prior on the motion or explicit priors for motion patterns that are likely to occur in neurophysiology data (e.g. discrete jumps and slow monotonic drift).
- Motion in 3 dimensions. In this work, we only considered motion in the depth direction along the laminar array, not horizontal motion in directions orthogonal to the array. While motion in depth is the most salient and detectable motion axis for laminar arrays, the motion function in MEDiCINe could be directly augmented to model 3-dimensional motion, which may offer benefits for users with 3-dimensional electrode arrays. This may also allow MEDiCINe to be used for motion estimation in recording modalities other than electrophysiology, such as calcium imaging.

**Figure 7.**
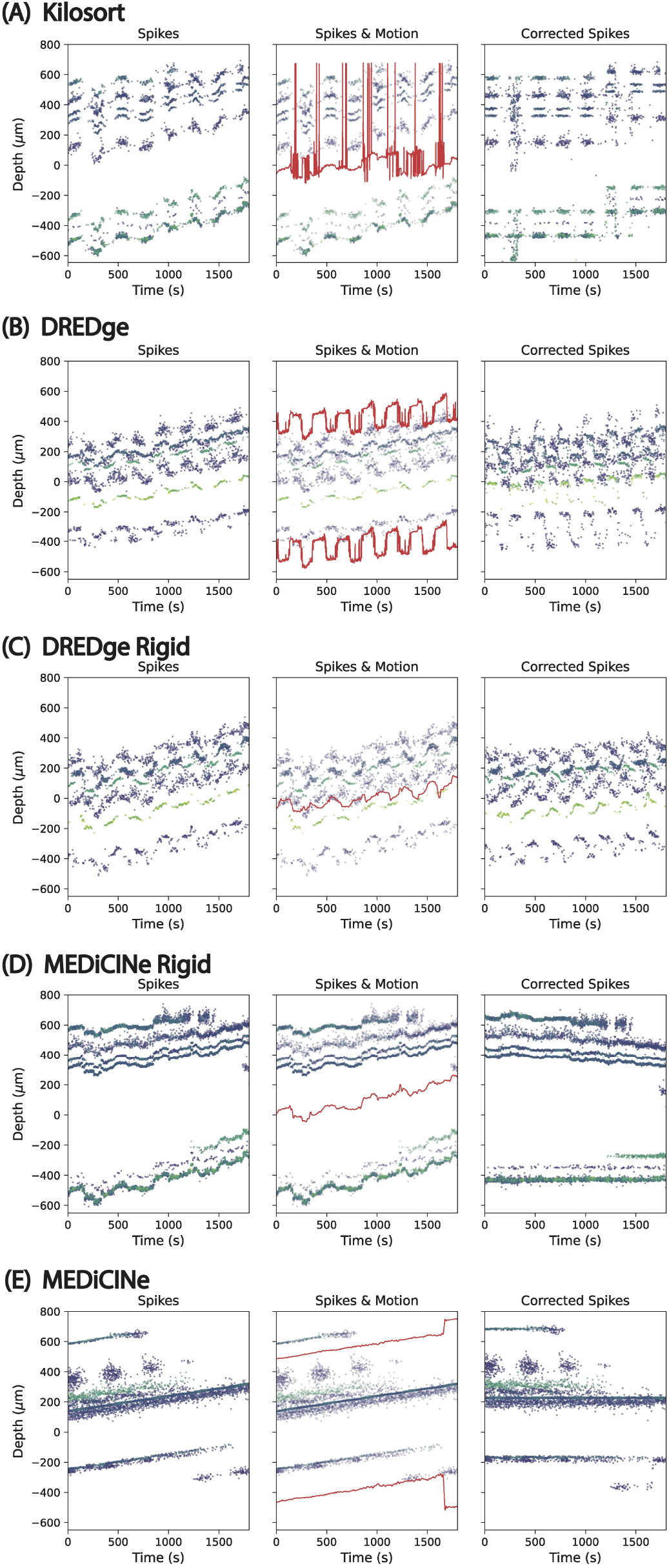
Failure Cases. **(A)** Kilosort motion estimation results for the simulated dataset for which the difference between Kilosort and the best method is greatest. This represent the worst failure case for Kilosort in our suite of simulated datasets. **(B) - (E)** Corresponding failure cases for the other methods.

We open-source the MEDiCINe model implementation and provide a website with documentation, demos, and instructions for installing and using MEDiCINe with just a few lines of code. We also open-source all code and data necessary for reproducing our results with instructions for how to do so.

## 5. Supplementary Material

### 5.1. Open-Sourcing and Reproducibility

To use MEDiCINe, please visit our MEDiCINe website https://jazlab.github.io/medicine. That website includes demos and instructions and for using MEDiCINe on your own data, including interfacing with Kilosort4 and SpikeInterface.

If you would like to reproduce the results in this work, please visit https://github.com/jazlab/medicine_paper for software, data, and instructions for reproducing the results in this manuscript.

### 5.2. MEDiCINe Details

#### 5.2.1. Motion Function

We parameterized the motion function by an array of size [*depth*_*bins, time*_*bins*]. For multiple depth bins, the depth bins uniformly divided the range from the deepest to the shallowest detected spike. We let *depth*_*bins* equal 2 for MEDiCINe. This allowed for non-uniform/non-rigid motion, was sufficiently flexible to capture motion well in all of our datasets, and agreed with the default depth discretization of DREDge on our simulated datasets. We let *depth*_*bins* equal 1 for the MEDiCINe Rigid model. We let *time*_*bins* equal the ceiling of the number of seconds in the dataset, allowing the model to capture motion at 1-second resolution. The model function also had a temporal smoothing kernel, for which we used a triangular kernel with 30-second support (15 seconds each for the ramp-up and ramp-down). We found this temporal resolution and smoothness to be sufficiently fine to capture motion well in all of our datasets

To compute the change in depth at a given time and depth, we computed the linear interpolation of the temporally smoothed motion array for that time and depth. We then applied a scaled hyperbolic tangent function to bound the motion by ±400*μm*.

#### 5.2.2. Activity Network

We parameterized the activity network by a multilinear perceptron with 14 input units, 2 fully connected hidden layers each with 256 units, and one output unit. The activation function in ReLU. The output was subject to a sigmoid function to force it to be a probability in [0, 1]. Given a depth and amplitude, to compute the probability of a corresponding spike we did the following:

- Normalize both depth and amplitude to lie in [0, 1], given the depths and amplitudes of all spikes in the dataset.
- Compute 6 depth features by taking sin (*x ∗*) *depth* for *x* in [1, 2, 4, 8, 16, 32]. Similarly, compute 6 amplitude features.
- Concatenate the depth and amplitude with their features into a 14-dimensional vector.
- Apply the MLP to this vector.

We added these sinusoidal features as inputs to the network because they helped optimization by allowing the MLP to more easily learn high-frequency modulations. In our experiments these features improved optimization convergence runtime by about a factor of 10.

#### 5.2.3. Depth Noise

To reduce the chance of converging to a local minimum, we added noise to the motion function output early in training. At the start of training this noise had standard deviation equal to 0.1 times the depth range of the data. This was linearly annealed to 0 throughout the first 2000 gradient steps of training.

#### 5.2.4. Optimization

We implemented the model in PyTorch and trained it with the Adam optimizer with learning rate 5 · 10^−4^ and gradient clipping of 1. We used batch size 8192, where each batch had 4096 spikes randomly sampled from the dataset and 4096 spikes randomly sampled from a uniform distribution with the same depth, amplitude, and time bounds as the spike dataset. We trained for 10,000 gradient steps. This took about 4 minutes on an M2 Macbook laptop, though we used a few CPUs on a computing cluster to run all 384 simulated datasets. We found that runtime could be improved by several factors with negligible impact on results by reducing the training steps to 5,000 and the batch size to 2048. GPU acceleration may further improve runtime, but we did not have a need for it.

### 5.3. Existing Methods

We used the following implementations for existing methods:

- Kilosort. We used the datashift function from the official Kilosort4 implementation, with the default hyper-parameters.
- DREDge. We used the official DREDge implementation in SpikeInterface version 0.101.2 with default parameters.
- DREDge Rigid. We used the “rigid_fast” method in SpikeInterface version 0.101.2. Note that this uses center-of-mass depth estimation instead of monopolar triangulation.

For details, see our repository for reproducing the results in this paper.

### 5.4. Simulated Data Results

#### 5.4.1. Dataset Generation

To generate simulated datasets with the parameters described in section 3.1, we used the MEArec library (Buccino and Einevoll, 2021). For each random component of motion and and instability we used a pseudorandom number generator with the same seed across datasets to ensure that our conditionalized results (Figure 4) were not influenced by random seed.

For the bimodal distribution of neuron density over depth, we sampled neuron depths from a mixture of two Gaussian distributions. Considering the length of the array to be the distance in depth between the deepest and the most shallow electrode, each of these Gaussian distributions had standard deviation equal to 10% of the array length, and their means were located at 15% and 85% of the array length.

Our firing rate stability statistics were generated as follows:

- Constant firing rates. The firing rate of each neuron was randomly uniformly sampled between 1Hz and 20Hz.
- Synchronous periodic firing rates. The firing rate of each neuron fluctuated with period 4 minutes as a truncated sine wave with shared phase, mean 1.5Hz, and truncated with a lower bound of 0.2 Hz.
- Asynchronous period firing rates. Same as synchronous periodic firing rates except each neuron’s phase was randomly and independently sampled.
- Ramping firing rates. Half of the neurons had constant firing rates randomly uniformly sampled between 1Hz and 20Hz. For the other half, each neuron was randomly attributed as either appearing or disappearing and given a random uniform time of appearance/disappearance. Its firing rate then ramped linearly in time between 0 at the appearance/disappearance time to a maximum randomly sampled between 1Hz and 20Hz at either the start (for disappearing) or the end (for appearing) of the session.

For depth-dependent motion, we let the motion linearly scale as a function of depth, with coefficient 1 for the deepest electrode and coefficient 0.5 for the most shallow electrode.

#### 5.4.2. Performance Quantification

To compute the mean absolute error of motion estimation in Figure 2-C, we first selected 11 depth levels evenly spaced from the deepest to the shallowest recorded spike. For each of these depth levels, we computed the ground truth motion through time, namely computing 11 of the red lines in Figure 2-A. Denote this by a matrix *M* of shape [ 11, *S*] where *S* is the number of seconds in the dataset and the values of *M* are ground truth Δ*depth* perturbations. The to compute the mean absolute error of a model on that dataset, we first computed the motion estimated by the model at those 11 depth levels, namely the depth perturbations predicted by the model. Denote this matrix 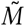 of shape [11, *S*]. We then compute the difference matrix 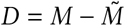. Then we center each row of *D* to have zero median, namely let *D* [*i*, :] ← *D* [*i*, :] − *median* (*D* [*i*, :]). We do this to avoid penalizing arbitrary biases in the estimated motion. Finally, we let the mean absolute error be the mean of the absolute values of all elements of *D*.

#### 5.4.3. Motion Estimation Results Breakdown

See Figure 4 for a breakdown of motion estimation results by parameter of the simulated data. This is a fine-grained look at the aggregate data in Figure 2. For each value of every parameter, MEDICINE and MEDICINE Uniform perform at least as well as existing methods, both in terms of mean absolute error and their ranking among methods.

#### 5.4.4. Benchmark Violin Plots

See Figure 5 for violin plot representations of the results in Figure 2. These show that the results do not appear to be due to outlier effects.

#### 5.4.5. Benchmark Method Rankings

In Figure 2-C and 2-D we showed mean absolute error of motion estimation and mean sorting accuracy for each method. These average results alone do not account for correlated variance across datasets that could cause, for example, a non-MEDiCINe method to out-perform MEDiCINe on the majority of the datasets. To eliminate this possibility, we consider for each simulated dataset the ranking of motion estimation methods (1 = best, 5 = worst), and show the mean ranking across all datasets for each method in Figure 6 for both mean absolute motion estimation error and relative sorting inaccuracy.

#### 5.4.6. Failure Cases

For each motion estimation method, we find the simulated dataset on which it performs worst relative to the best method. See Figure 7 for plots of the results for these worst-case datasets. These give an intuition for the conditions under which each method may fail and the manner of failure.

#### 5.4.7. Spike Sorting Accuracy

To evaluate the performance of spike sorting after motion correction by a model, we first corrected the raw recording by the estimated motion using Kriging interpolation, then ran Kilosort4 spike sorting. Denote the ground truth neurons by {*g*_*i*_} _1 ≤ *i* ≤ *N*_ and the sorting output units by {*s*_*j*_} _1 ≤ *j* ≤ *M*_. For each pair of ground truth neuron *i* and sorting unit *j* we computed the accuracy *Accuracy*_*i,j*_ as

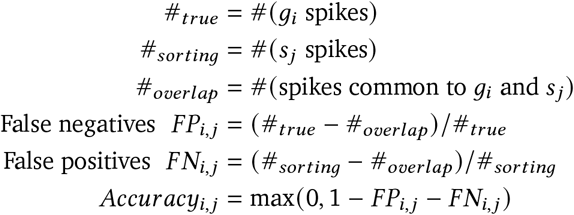

where a spikes in common to *g*_*i*_ and *s*_*j*_ were all spikes in *s*_*i*_ that occurred within 1ms of a spike in *g*_*i*_. For each ground truth neuron *g*_*i*_ we let *Accuracy*_*i*_ = max_1≤ *j* ≤ *M*_ *Accuracy*_*i,j*_. This gives us an accuracy measure in [0, 1] for each ground truth neuron. Accuracy computed in this way is a standard measure of spike sorting accuracy (Garcia et al., 2024; Pachitariu et al., 2024). To reduce this to a single score we let the sorting inaccuracy be *Inaccuracy* =Σ_1≤*i* ≤*N*_ (1 − *Accuracy*_*i*_,_*j*_) Given spike sorting inaccuracies for all motion estimators on a particular dataset, we compute the relative inaccuracy of estimator *A* as

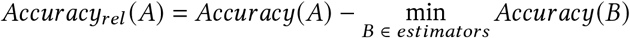

In Figure 2-D we showed the average relative inaccuracy for each method across these datasets. In Figure 8 we show the accuracy plots for each method on each of the 40 datasets for which we ran spike sorting. These plots show accuracy as a function of unit number (sorted by accuracy) in the same style as (Garcia et al., 2024) and (Pachitariu et al., 2024).

**Figure 8.**
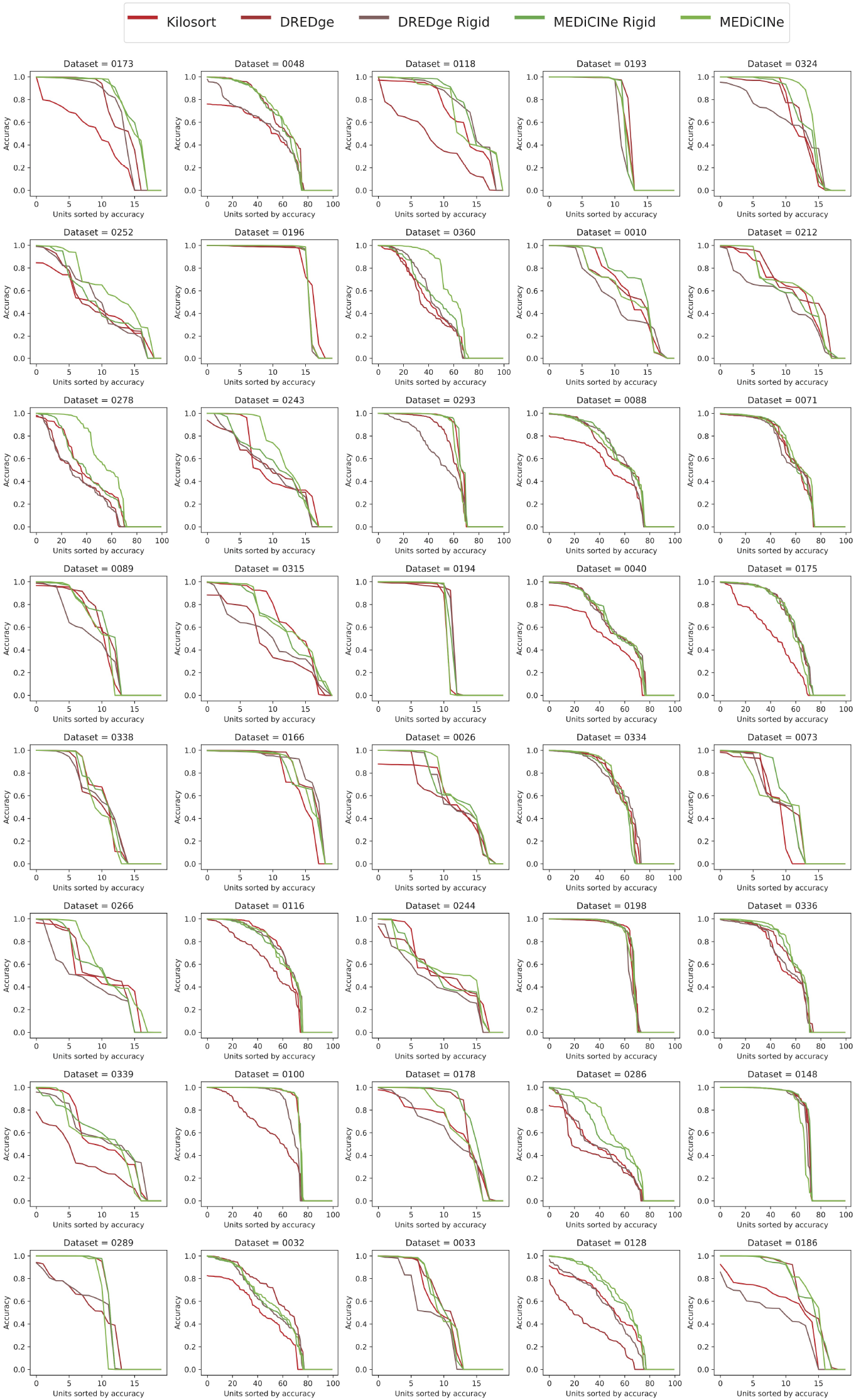
Spike Sorting Accuracy. Accuracy as a function of unit (sorted by accuracy) for Kilosort4 sorting results for each motion estimation method on each of the 40 datasets for which we ran spike sorting.

**Figure 9.**
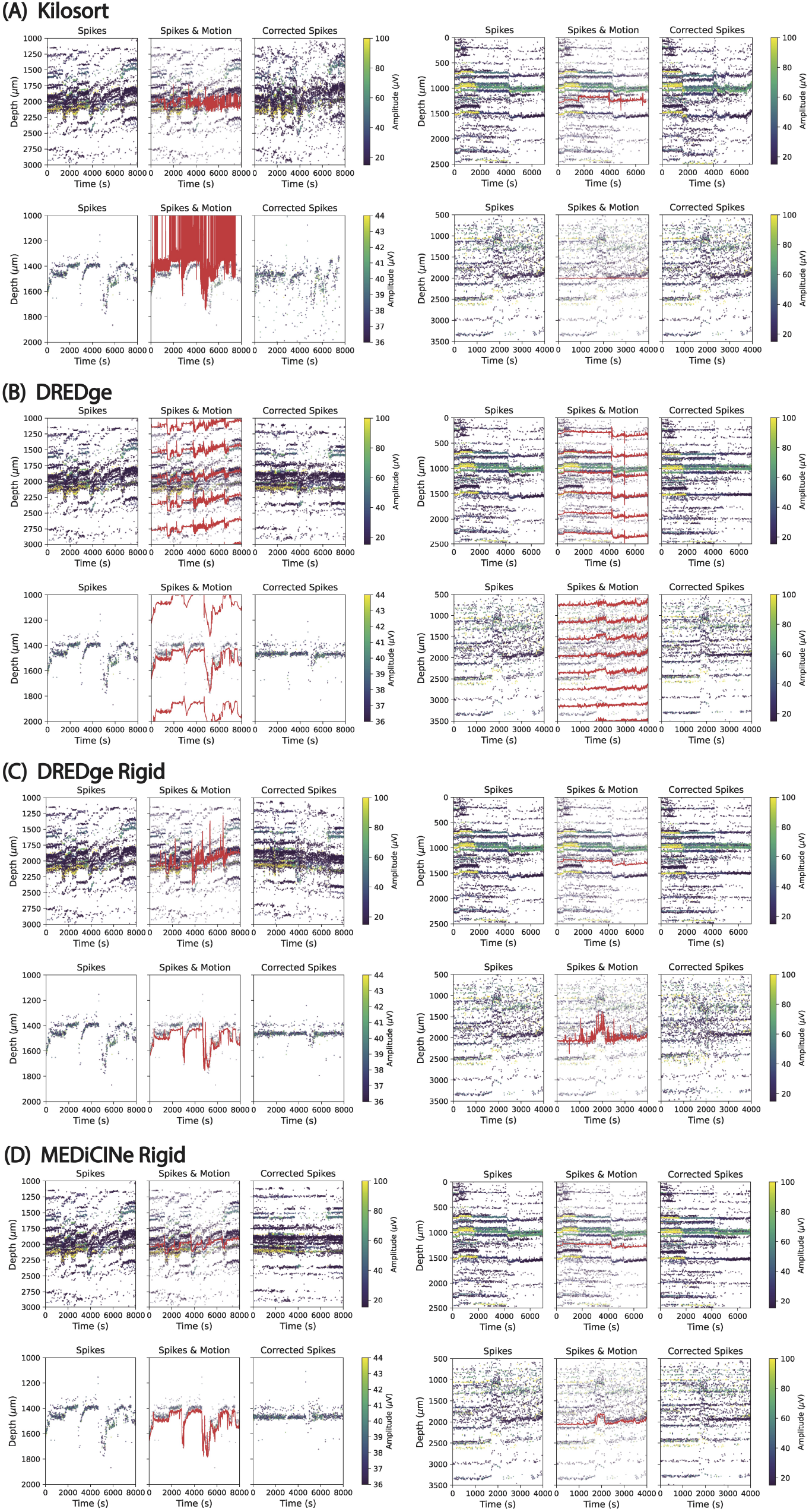
Non-MEDICINE Results for NHP Datasets. This shows the results for all Non-MEDICINE methods for each of the NHP datasets shown in Figure 3.

### 5.5. Primate Data Results

See Figure 9 for results from all non-MEDICINE models on the nonhuman primate datasets shown in Figure 3. Comparing these to the MEDICINE results in Figure 3, by eye MEDICINE looks more aligned with the underlying motion than the other methods. The spike data in these panels was thresholded to remove very small spikes for visualization purposes only, but the methods were run on the monopolar triangulation spike detection output directly from SpikeInterface.

